# *In vitro* reconstitution of the *M.tb* proteasome core particle reveals conserved aspects of bacterial proteasome assembly

**DOI:** 10.1101/2023.07.05.547829

**Authors:** Anupama Kante, Jeroen Roelofs, Eric J. Deeds

## Abstract

According to the WHO, one in three people in the world has a latent tuberculosis infection. Tuberculosis is caused by the bacterium *Mycobacterium tuberculosis (M.tb).* The development of multi-drug resistant (MDR) tuberculosis indicates a need for novel treatments. Hence, it is important to find a second line of treatment for patients infected with MDR tuberculosis. The proteasome is known to be necessary for survival under stress and pathogenicity in *M.tb*. However, our ability to use the proteasome as drug target has been limited by our abilities to screen for inhibitor compounds *in vitro.* The proteasome is a protease complex that degrades proteins and is crucial for the maintenance of protein homeostasis within cells. Like many protein complexes, the proteasome must assemble into a specific quaternary structure in order to be active. Specifically, the proteolytically-active proteasome Core Particle (CP) consists of 28 subunits (14 α and 14 β) that must assemble into a barrel-like structure in order to become catalytically active. Hence, understanding the assembly process in not only important from a basic cell biological perspective, but may also serve as the basis for the discovery of novel assembly inhibitors. In this study, we have established for the first time a protocol to express and purify the *M.tb* α and β subunits separately *in vitro.* The subunits are soluble monomers on purification and only assemble into active CPs upon reconstitution. Our assembly experiments revealed that *M.tb* CP assembly pathway is almost certainly identical to that seen in previous experiments on the CP from the bacterium *Rhodococcus erythropolis* (*R.e*), but assembly in *M.tb* is much slower. Interestingly, we found that subunits from *M.tb* and *R.e* spontaneously self-assembled into active hybrid proteasomes on reconstitution with each other, despite having only 65% sequence similarity. Our work thus strongly suggests that the CP assembly pathway is conserved across bacteria, and the ability to perform *in vitro* assembly experiments on the *M.tb* proteasome opens up the possibility of performing critical experiments, including screening for potential molecules that could inhibit assembly, directly in this clinically-relevant organism.

## Introduction

Tuberculosis is a disease with a high mortality rate. It is caused by the actinomycete bacterium *Mycobacterium tuberculosis* (*M.tb*) (1). While there are antibiotics that are effective in treating tuberculosis infections, antibiotic-resistant *M.tb* strains have evolved, and their growing prevalence poses an increasingly serious public health problem (1). *M. tuberculosis* strains have used compensatory evolution, epistasis, clonal interference, cell wall integrity, efflux pumps, and target mimicry to develop resistance (1). It is thus critical to understand the biological processes in *M.tb* that contribute to pathogenesis in molecular detail, so those processes can be targeted in order to develop novel antibiotics that are effective against strains that are resistant to currently-available antibiotics.

*M.tb* infects macrophages in the host (2). The macrophages use the Nitric Oxide Synthase 2 (NOS2) enzyme, which generates the highly reactive free radical nitric oxide (NO), as part of their response to an active infection (3). When the immune response is non-sterilizing (in other words, unable to completely clear the bacteria), the resulting persistent infections can lead to active tuberculosis in the future (2). Transposon mutagenesis screens for hyper-susceptibility to NO in *M.tb* identified both *PrcA* and *PrcB*, the genes that code for the α and β proteasome Core Particle (CP) subunits in *M.tb*, suggesting the proteasome is critical for survival of the bacteria under nitrosative stress (4). Besides the two proteasome subunits this screen also identified the *Rv2115c* gene, which encodes an AAA-ATPase (5). Interestingly, a peptidyl boronate proteasome inhibitor reduced growth of *M.tb*, and mutation of the *Rv2115c* gene reduced fitness of the bacteria and its ability to cause infection and kill the host(6). These findings indicate that the proteasome is essential for survival and pathogenicity of *M.tb* in infected hosts.

Most bacteria express two ATP-dependent proteolytic complexes HslUV and Lon (7). Mycobacterium tuberculosis belongs to the phylum Actinomycetes, which are the only eubacteria to possess a proteasome. *M.tb* proteasomes have been purified as an assembled complex by co-expressing the *PrcA* and *PrcB* genes in E*.coli* (*2*). However, it has been impossible to express and purify the *PrcA* and *PrcB* proteins separately due to their insolubility, making it impossible to study the assembly process *in vitro.* This is in contrast to the CP from the related actinomycete bacterium *Rhodococcus erythropolis* (*R*.e), where these two proteins can be readily expressed separately as monomers(8) (9–11). Negative stain electron microscopy (EM) images of *M.tb* proteasomes show a conserved four stacked ring structure similar to the 20S proteasome from *R.e,* the archaeon *Thermoplasma acidophilum,* yeast and humans (2). Negative stain EM studies also showed that the TB proteasome exists in two conformations—open gate and closed gate (2). Substrate entry into the catalytic core and accessibility to the proteolytic site is only possible in the open gate conformation of the alpha subunit (2). The *M.tb* proteasomes were shown to be enzymatically active with a preference towards the flurogenic substrate Z-VLR-AMC (2). Known inhibitors of eukaryotic proteasome reduced the proteolytic activity of *M.tb* proteasomes(2).

The *M.tb* proteasome is a very promising target for novel anti-tuberculosis drugs. While screens have identified possible enzymatic inhibitors of the *M.tb* proteasome, the development of an *M.tb*-specific inhibitor has been challenging, and to our knowledge no molecule targeting the *M.tb* proteasome is currently in clinical trials for the treatment of TB. Targeting the assembly process could represent a novel and unique way to reduce *M.tb* proteasome activity. While structural studies have provided detailed information about the fully assembled CP in *M.tb*, very little is known about the assembly process. *In vitro* self-assembly studies with the *R.e* proteasome have led to the suggestion that bacterial proteasomes assemble via an “α-β” dimer pathway (12). In this pathway, seven α-β hetero-dimers form a Half Proteasome (HP) and two HPs dimerize to form the CP. But, this pathway remains untested in tuberculosis due to the inability to reconstitute *M.tb* proteasomes *in vitro*.

Here, we have expressed and purified the α and β subunits from *M.tb* separately, characterized them and demonstrated spontaneous *in vitro* assembly into active proteasome CP complexes when mixed together. Assembly time courses resolved on native gels revealed clear HP and CP bands, similar to those observed with the *R.e* subunits. In *R.e*, there is a distinct separation of time scales between HP formation (which happens extremely rapidly) and HP dimerization, which under standard experimental conditions can take one or two hours to proceed to completion (11, 12). We found a similar separation of timescales in the *M.tb* proteasome, but under similar conditions dimerization was much slower, only going to completion over the course of 12-24 hours. We also demonstrated that subunits from *M.tb* and *R.e* were compatible with each other; on mixing *M.tb* α with *R.e* β and *vice-versa* they assembled into active CPs. Finally, increasing the concentration of the subunits produced a characteristic relationship between subunit concentration and assembly efficiency, essentially identical to that observed for the *R.e* subunits. Specifically, we found that the two CPs exhibit essentially the same level of kinetic trapping (13).

Taken together, these findings strongly indicate that the assembly pathway is conserved between *M.tb* and *R.e* and likely all bacteria. Both self-assembly experiments and MD simulations of the HP from *R.e* suggest that a flexible region of the β subunit (specifically, a part of the β propeptide that is cleaved off during autocatalytic activation of the CP) is responsible for regulating the timescale of HP dimerization (8, 11). It is unclear if changes to the sequence of this region in *M.tb* are responsible for the dramatic decrease in the rate of HP dimerization in this case. Our work also demonstrates that kinetic trapping is conserved between *R.e* and *M.tb*, underscoring the importance that this phenomenon has likely played in the evolution of assembly pathways of the proteasome and other macromolecular machines (13). Finally, the ability to spontaneously assemble *M.tb* proteasomes from its subunits *in vitro* will facilitate future detailed *in vitro* studies of *M.tb* proteasome assembly, *in vitro* enzymatic characterization, structural studies of assembly intermediates, and screens for specific CP assembly inhibitors.

## Materials & methods

### Cloning the PrcA and PrcB gene

The open reading frame (ORF) for *M.tb PrcA* (we used an open gate mutant lacking the first 8 codons) and *PrcB* genes (full length) were optimized for expression in *E.coli* and synthesized (see supplement) in a pUC57 vector by Genscript. Gene sequences are listed in the Supplement. The *PrcA* and *PrcB* genes were sub-cloned in the pET His6 TEV LIC cloning vector (2B-T, a generous gift from the Scott Gardia lab) (Addgene# 29666) and pET His6 TEV LIC cloning vector (2A-T, a generous gift from the Scott Gardia lab) (Addgene # 29665) using Ligation Independent Cloning to generate the pAKTB-α and pAKTB-β plasmids, respectively. The ORFs were PCR amplified using the primers listed in Sup Table 1, with LIC tags prescribed by the manufacturer. PCR products were treated with T4 polymerase in presence of the relevant dNTPs (Supp Table 1). The 2B-T and 2A-T plasmids were linearized using the restriction enzymes Sspl and EcoRV respectively. The linearized plasmids were treated with T4 polymerase in presence of the dNTPs recommended by the manufacturer. The insert and the vector were annealed, and the resultant plasmids were transformed in E.Coli BL21-DE3 cells (NEB).

### Protein expression and purification

Plasmids containing the α and β genes were freshly transformed in E.Coli BL21-DE3 cells (NEB). Transformed cells were plated on LB agar plates containing 100ug/ml Carbenicillin and grown overnight at 37^0^C overnight. A single colony of from the plate was inoculated in 100ml of Luria-Bertani (LB) broth containing 100ug/ml Carbenicillin and incubated under shaking conditions of 200 rpm at 37^0^C overnight. 50 ml of the overnight starter culture was used to inoculate per 1L of LB media (total 4L media was used to grow the cells). Cells were allowed to grow up to OD_600_ of 0.6 and induced with 1 mM IPTG. Cells containing the α subunit were induced at 15^0^C, overnight whereas cells containing the β subunit were induced at 30^0^C for 4hours. Cells were harvested by centrifugation at 4000 rpm (JS-4.750 rotor in a Avanti-J15R centrifuge) for 15 mins at 4^0^C. Cell pellets were resuspended in 50ml binding buffer (specified below for each subunit) and stored at -80^0^C. Next, cells were thawed, and lysed by sonication at 45% amplitude for 4 mins with a 2 sec on and 20 sec off cycle using a Fischer Scientific Sonic dismembrator model 500. Lysate was cleared by centrifugation at 10,000 rpm (JA-10.100 rotor in a Avanti-J15R centrifuge) for 30 mins at 4^0^C. Proteins were purified from the clarified lysate using a GE AKTApure FPLC system using immobilized metal affinity chromatography. The β subunit was further purified using anion exchange chromatography.

### Purification of the α subunit

In the pAKTB-α plasmid, the PrcA gene is flanked by a N-terminal TEV cleavable 6-His tag. In the first step of purification, the clarified lysate was loaded on a Nickel affinity column (GE His-trap 5ml column), the column was then washed with 10 column volumes (CV) of binding buffer (50mM Tris-HCL, 500mM NaCl, 5mM DTT, pH 8.0 at 4^0^C). The bound protein was eluted with a linear gradient from 0 to 100% of the elution buffer (50mM Tris-HCL, 500mM NaCl, 5mM DTT, 500mM Imidazole, pH 8.0 at 4C) over 20CV. Peak fractions were analyzed using SDS-PAGE. Fractions containing the α subunit were pooled and incubated with 1:50 volumetric ratio of 62μM TEV protease (1 ml of TEV for 50 ml of protein fraction) and dialyzed in the same binding buffer overnight at 4^0^C. Next day, the TEV cleaved protein was again loaded on Nickel affinity column (GE His-trap 5ml column). The cleaved protein was separated from the His tag and the TEV protease using the protocol described above. The TEV-cleaved protein does not bind the column and is hence collected in the flow through fractions. Fractions containing the protein were analyzed using SDS-PAGE. 20% v/v glycerol was added to the pooled fractions and aliquotes of 1 ml were frozen in liquid nitrogen and stored at -80^0^C.

### Purification of β subunit

In the pAKTB-β plasmid, the Prc B gene is flanked by a C-terminal 6-His tag. In the first step of the purification, the cleared lysate was loaded on a Nickel affinity column (GE His-trap 5ml column), the column was then washed with 10 column volumes (CV) of binding buffer (50mM Tris-HCL, 500mM NaCl, pH 8.0 at 4^0^C). The bound protein was then eluted with a linear gradient (0-100%B) of the elution buffer (50mM Tris-HCL, 500mM NaCl, 500mM Imidazole, pH 8.0 at 4^0^C) over 20CV. Peak fractions were analyzed using SDS-PAGE. Fraction containing the protein were dialyzed in anion exchange binding buffer (20mM Tris-HCL, 20mM NaCl, pH 8.0 at 4^0^C) for approximately 16h at 4^0^C. The dialyzed protein was then purified using a GE 5ml Hi-trap anion exchange column. Unbound protein and contaminants were washed with 10 CV of binding buffer (20mM Tris-HCL, 20mM NaCl, pH 8.0 at 4^0^C). Bound protein was eluted using a linear gradient (0-100% B) of the elution buffer (20mM Tris-HCL, 1M NaCl, pH 8.0 at 4^0^C) over 20CV. Peak fractions were analyzed using SDS-PAGE. Fractions containing the protein were pooled together. 20% v/v glycerol was added to the pooled fractions and aliquotes of 300µL were frozen in liquid nitrogen and stored at -80^0^C.

### Dynamic Light scattering

Each subunit protein was thawed on ice. The subunits were buffer exchanged into HNE (20mM HEPES, 300mM, 1mM EDTA, pH 7.0) using Millipore!s Amicon Ultra-15, Ultracel-10K Centrifugal filter units. Protein concentrations were estimated by measuring absorbance at 280nm, using the molar extinction co-efficient of 16390 for *M.tb* α and 24410 for M.tb β. A final solution of 8µM of each subunit was prepared in HNE. Protein samples were analyzed using a Wyatt Plate Reader II. Dispersant was set to water and data was acquired at 20^0^C.

### In vitro assays and reconstitution

The subunits were buffer exchanged into HNE (20mM HEPES, 300mM, 1mM EDTA, pH 7.0) using Millipore!s Amicon Ultra-15, Ultracel-10K Centrifugal filter units. Protein concentrations were estimated by measuring absorbance at 280nm, using the molar extinction co-efficient of 16390 for *M.tb* α and 24410 for M.tb β. A final solution of 8µM of each subunit was prepared in HNE. To determine temperature stability of subunits, the individual subunits were incubated at 4^0^C, 10^0^C, 20^0^C, 30^0^C, 37^0^C and 42^0^C for 1 hour. For assembly assays, equal volume of 8µM α and 8µM β were incubated together at 30^0^C for 48hrs. For time courses of assembly, equal volume of 8µM α and 8µM β were incubated together at 30^0^C for 120h (5 days), 96 h (4 days), 72h (3 days), 48h (2 days), 36h, 24h, 12 h, 6 h, 3 h, 1.5 h, 45 mins and 0 mins. Equal volume of loading dye (0.8M HEPES,0.1% Bromophenol Blue, 20% Glycerol) was added to the proteins. 20µL of sample was applied on an Invitrogen Novex ™ Wedge well ™ 4-20% Tris-Glycine gel. Gel was run for 250mins, 120Volts at 4^0^C. Gel was stained with Sypro ruby protein stain (Thermofischer) as described by the manufacturer, imaged using a proprietary Sypro ruby imaging protocol in Biorad ChemiDoc imager and visualized using Imagelab software.

### Cross Assembly

The α and β subunits from *Rhodococcus(R.e)* were expressed and purified as described in the supplement. The α and β from both *M.tb* and *R.e* were buffer exchanged into HNE (20mM HEPES, 300mM, 1mM EDTA, pH 7.0) using Millipore!s Amicon Ultra-15, Ultracel-10K Centrifugal filter units. Protein concentrations were estimated by measuring absorbance at 280nm and using the molar extinction co-efficient of 16390 for *M.tb* α and *R.e* α, 17880 for R.e β and 24410 for M.tb β. A final solution of 4µM of each subunit was prepared in HNE. Equal volumes of *M.tb* α and β were mixed together as the first control. Equal volumes of *R.e* α and β were mixed together as the second control. The cross assembly reactions composed of the following ratios of subunits: *M.tb* α + *R.e* β, *R.e* α + *M.tb* β, 0.5 *M.tb* α + 0.5 *R.*e α + *R.e* β, 0.5 *M.tb* α + 0.5 *R.e* α + *M.tb* β, *R.e* α + 0.5 *R.e* β + 0.5 *M.tb* β, *M.tb* α + 0.5 *R.e* β + 0.5 *M.tb* β. All assembly reactions were incubated for 24 hours at 30^0^C. At the end of 24hrs, equal volume of loading dye (0.8M HEPES,0.1% Bromophenol Blue, 20% Glycerol) was added to the assembly reactions. Samples were the applied on a Invitrogen Novex ™ Wedge well ™ 4-20% Tris-Glycine gel. Gel was run for 8 hours, 120Volts at 4^0^C. Native gel was stained with Sypro ruby protein stain as described by the manufacturer, imaged using Biorad ChemiDoc imager and visualized using Imagelab software.

### Activity Assay

Samples from the assembly and cross assembly assays were loaded onto two gels. One gel was used to visualize the protein bands using the Sypro Ruby stain and the second gel was used to perform an activity assay. For the latter, gels were rinsed in water for 10 mins and then transferred into activity assay buffer containing HNE (20mM HEPES, 300mM, 1mM EDTA, pH 7.0) and 50mM of the fluorogenic substrate Z-VLR AMC (Adipogen Life sciences). The gel was incubated in the activity assay buffer for 20 mins at 37^0^C with intermittent shaking. The gel was imaged using Biorad ChemiDoc imager using UV transillumination (302nm excitation) light source and a standard filter (590/110 nm). Imagelab software was used for visualizing and processing the gels.

### Effect of subunit concentration on Assembly

The α and β subunits were transferred in to HNE buffer (20mM HEPES, 100mM NaCl,1mM EDTA, 5mM DTT PH 7.0) and concentrated separately using Amicon Ultra 10K centrifugal filters (Millipore). Protein concentrations were estimated to be roughly 35µM by measuring absorbance at 280nm and using the molar extinction co-efficient of 16390 for α and 24410 for β. These stocks were then diluted to 0.5µM, 1µM, 2µM, 4µM, 8µM, 16µM and 32µM using the HNE buffer. Both subunits were mixed in equal volumes to obtain a final subunit concentration of 0.25 µM, 0.5 µM, 1 µM, 2 µM, 4 µM, 8 µM and 16 µM. The assembly reaction was allowed to proceed for 5 days at 30°C with intermittent spinning in a micro-centrifuge to pull down the condensation. Assembly reactions were mixed with equal volume of loading dye (0.8M HEPES,0.1% Bromophenol Blue, 20% Glycerol). Invitrogen Novex ™ Wedge well ™ 4-20% Tris-Glycine gel. The gel was run for 12 hours, 120V at 4^0^C. Native gel was stained with Sypro ruby protein stain as described by the manufacturer, imaged using Biorad ChemiDoc imager and visualized using Imagelab software.

## Results

### Purification and characterization of M.tb proteasome subunits

Previous efforts to study *M.tb* proteasome assembly *in vitro* have been unsuccessful due to an inability to purify the individual subunits. Here, we created new constructs for heterologous expression of each subunit (**Fig. 1A**). The M.tb CP exists in two states - a closed gate and an open gate. The proteolytic site is accessible to the substrate only in the open gate confirmation. The first 8 amino acid residues of the α subunit have shown to be involved in transition between the two states. Deletion of these 8 amino acids increased the activity of the *M.tb* CP to 14 fold (2). Hence, we deleted these residues in the α subunit in order to produce active particles that can be used to perform future *in vitro* drug screens. Both the α and β subunits were purified under native conditions and the N-terminal His tag of the α protein was removed using TEV protease. **Fig. 1B** shows protein profile during the different steps in the α protein purification protocol. **Fig. 1C** shows the protein profile during the different steps in the β protein purification protocol. Both proteins behaved well in solution and no precipitation was observed even at higher concentrations. We speculate that the location of the His-Tags on both the constructs improved their solubility. In addition, the deletion of the first 8 amino acids of the α subunit may have also improved the solubility of the protein.

**Fig 1:**
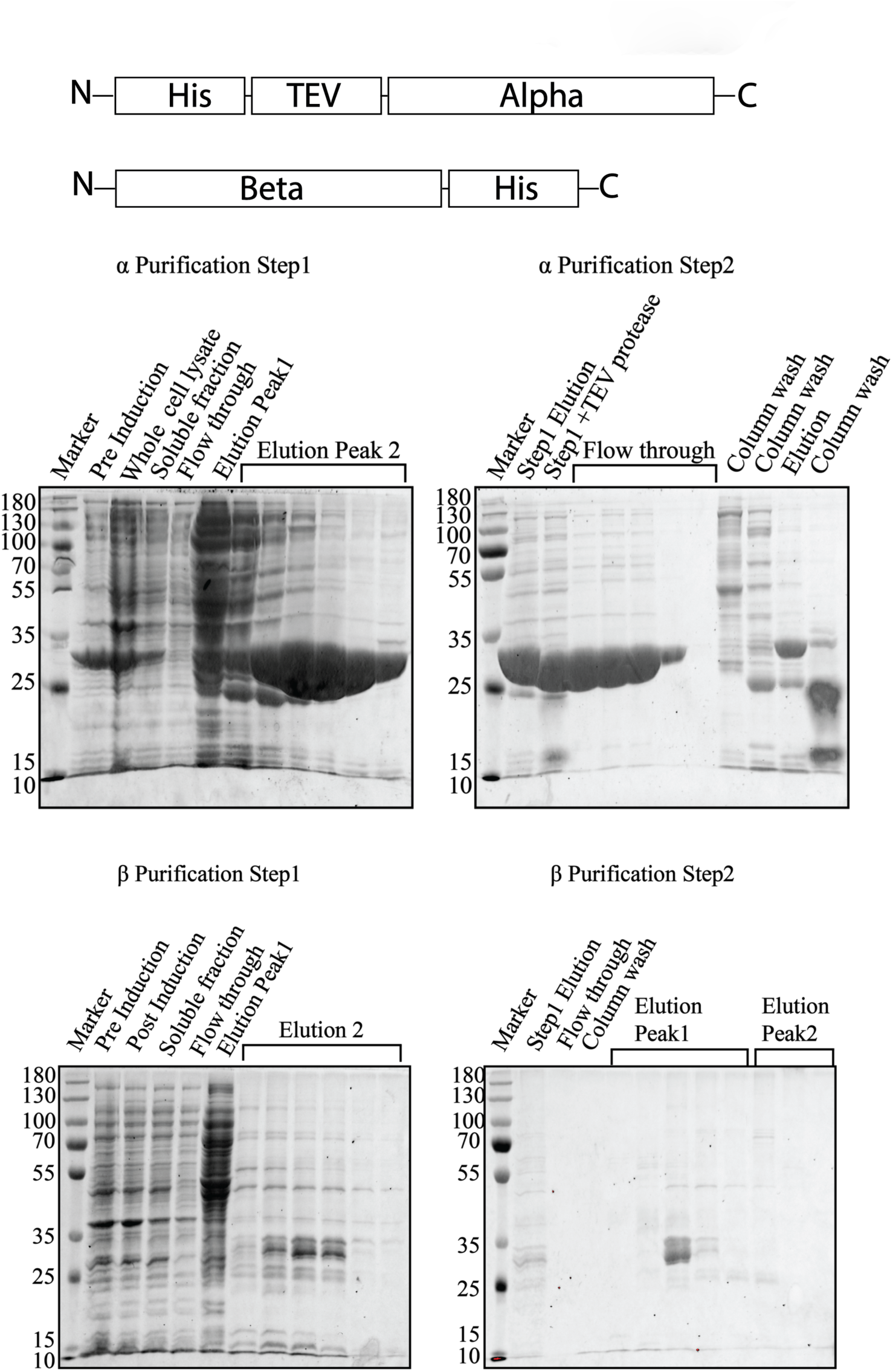
Expression and purification of M.tb proteasome subunits. **(A)** Schematic of the constructs used to purify the *M.tb* α and β subunits. **(B)** SDS-PAGE gels showing the purification profile of M.tb α. The results of the first step of purification are shown on the left, and the results of the second step on the right. **(C)** SDS-PAGE gels showing the purification profile of the *M.tb* β subunit. As in (B), the first step is shown to the left and the second step to the right.

To assess whether the soluble, purified subunits were homogenous monomers or were forming oligomers, we performed dynamic light scattering studies of individual subunits under native conditions. Single peaks were observed for each subunit (**Fig. 2A** and **2B**). This indicated the solutions composed of homogeneous protein species. Next, the subunits were mixed together in equimolar ratio and allowed to assemble for 24 hours at 30^0^C Fig 1E. Dynamic light scattering of these samples indicated that the average molecular weight of complexes in the assembly reaction was 490.7 kDA, which corresponds to the size of the HP. The observed values of Hydrodynamic radius, percent intensity and percent number are summarized in **Table 1**. Taken together, this indicates that it is possible to purify individual α and β subunits that are monomeric in solution and that they have the potential to form larger complexes when combined.

**Fig 2:**
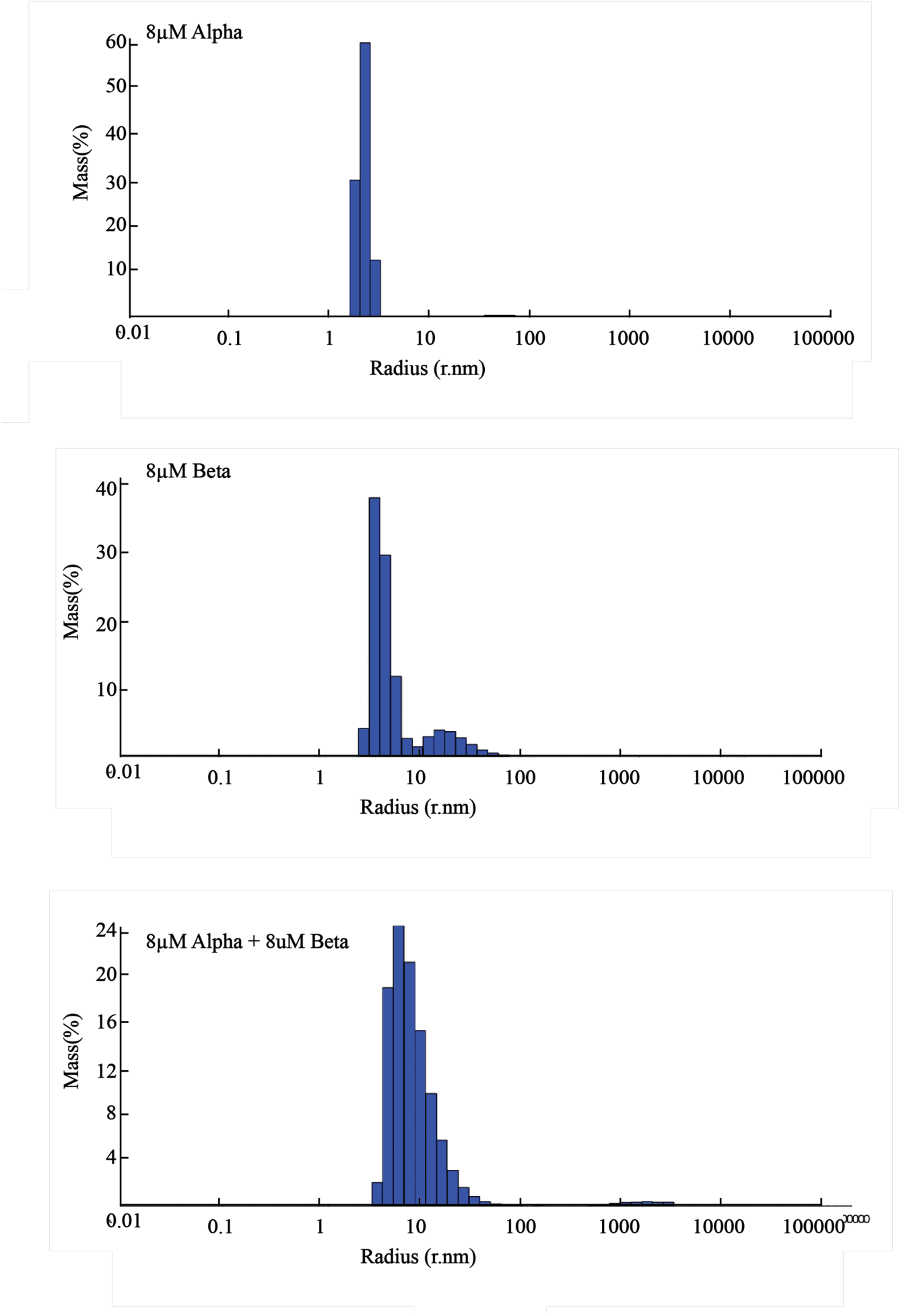
Dynamic Light scattering analysis of the individual M.tb subunits and their reconstitution product. **(A)** Distribution of observed hydrodynamic radiuses in a solution of 8µM α. **(B)** Distribution of observed hydrodynamic radiuses in a solution of 8µM β. **(C)** distribution of observed hydrodynamic radiuses in a reconstitution reaction made up of 4µM α and 4µM β.

**Table 1:**
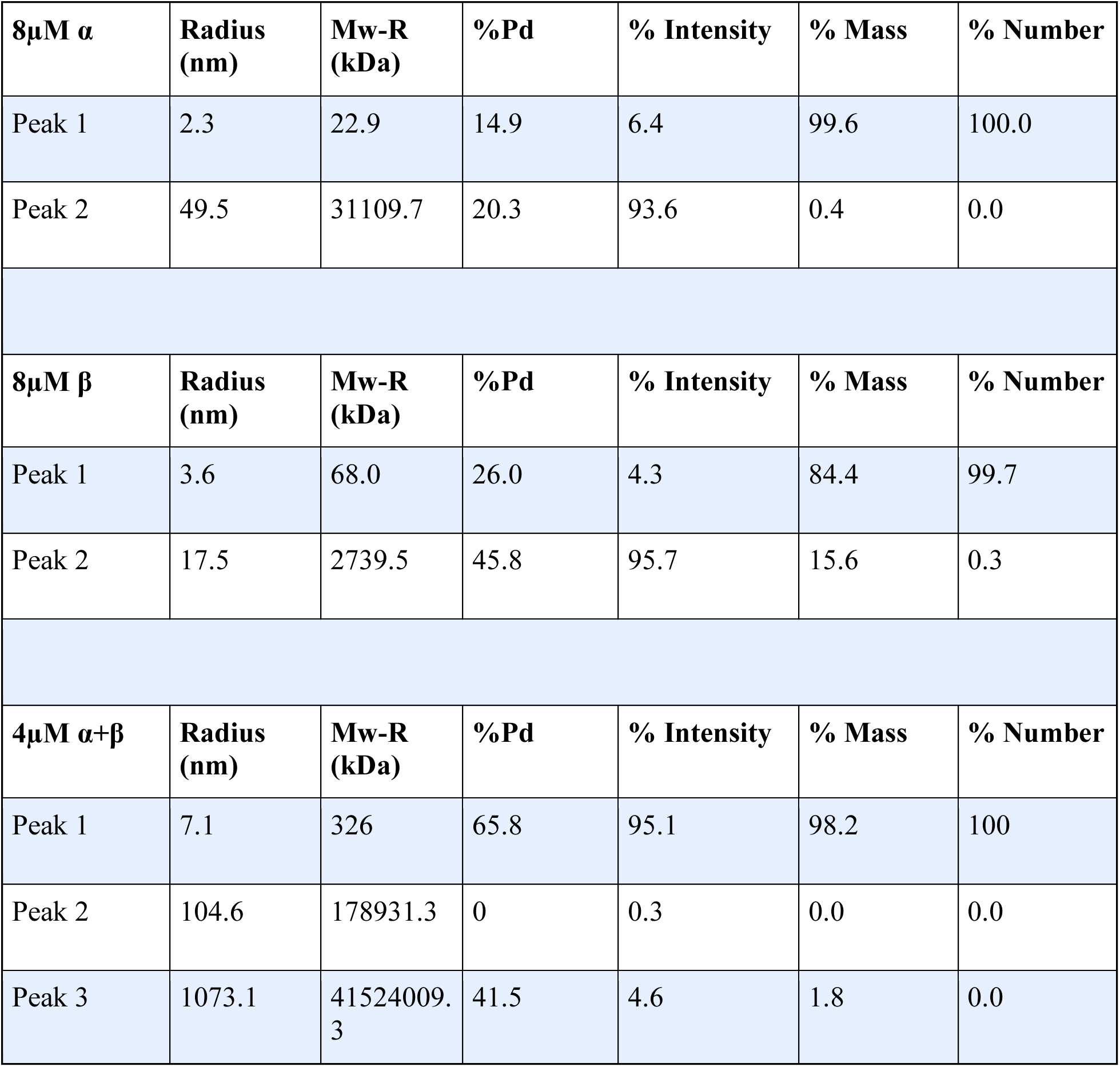
Summary table of DLS.

| **8μM $\alpha$** | **Radius**<br>**(nm)** | **Mw-R**<br>**(kDa)** | **%Pd** | **% Intensity** | **% Mass** | **% Number** |
| --- | --- | --- | --- | --- | --- | --- |
| Peak 1 | 2.3 | 22.9 | 14.9 | 6.4 | 99.6 | 100.0 |
| Peak 2 | 49.5 | 31109.7 | 20.3 | 93.6 | 0.4 | 0.0 |
| **8μM $\beta$** | **Radius**<br>**(nm)** | **Mw-R**<br>**(kDa)** | **%Pd** | **% Intensity** | **% Mass** | **% Number** |
| Peak 1 | 3.6 | 68.0 | 26.0 | 4.3 | 84.4 | 99.7 |
| Peak 2 | 17.5 | 2739.5 | 45.8 | 95.7 | 15.6 | 0.3 |
| **4μM $\alpha+\beta$** | **Radius**<br>**(nm)** | **Mw-R**<br>**(kDa)** | **%Pd** | **% Intensity** | **% Mass** | **% Number** |
| Peak 1 | 7.1 | 326 | 65.8 | 95.1 | 98.2 | 100 |
| Peak 2 | 104.6 | 178931.3 | 0 | 0.3 | 0.0 | 0.0 |
| Peak 3 | 1073.1 | 41524009.3 | 41.5 | 4.6 | 1.8 | 0.0 |

However, to conduct a comprehensive analyses of the M.tb proteasome assembly under different biophysical conditions, we had to first test if the subunits are stable at the different temperatures. For this we incubated the subunits at increasing temperatures and analyzed the resulting solutions by native gel PAGE to determine stability and tendency to aggregate (*Fig. 3*). The amount of α subunit migrating at the expected molecular weight remained the same from 4^0^C to 37^0^C and showed a slight reduction at 42^0^C, which was accompanied by an increased protein stain at the top of the gel, suggesting some modest aggregation at 42^0^C. The β subunit was stable up to 30^0^C and displayed some aggregation at 37^0^C and 42^0^C as seen by the smearing pattern. In all, both proteins were stable up to 30^0^C under these experimental conditions, and we used a temperature of 30^0^C to perform all subsequent self-assembly experiments.

**Fig 3:**
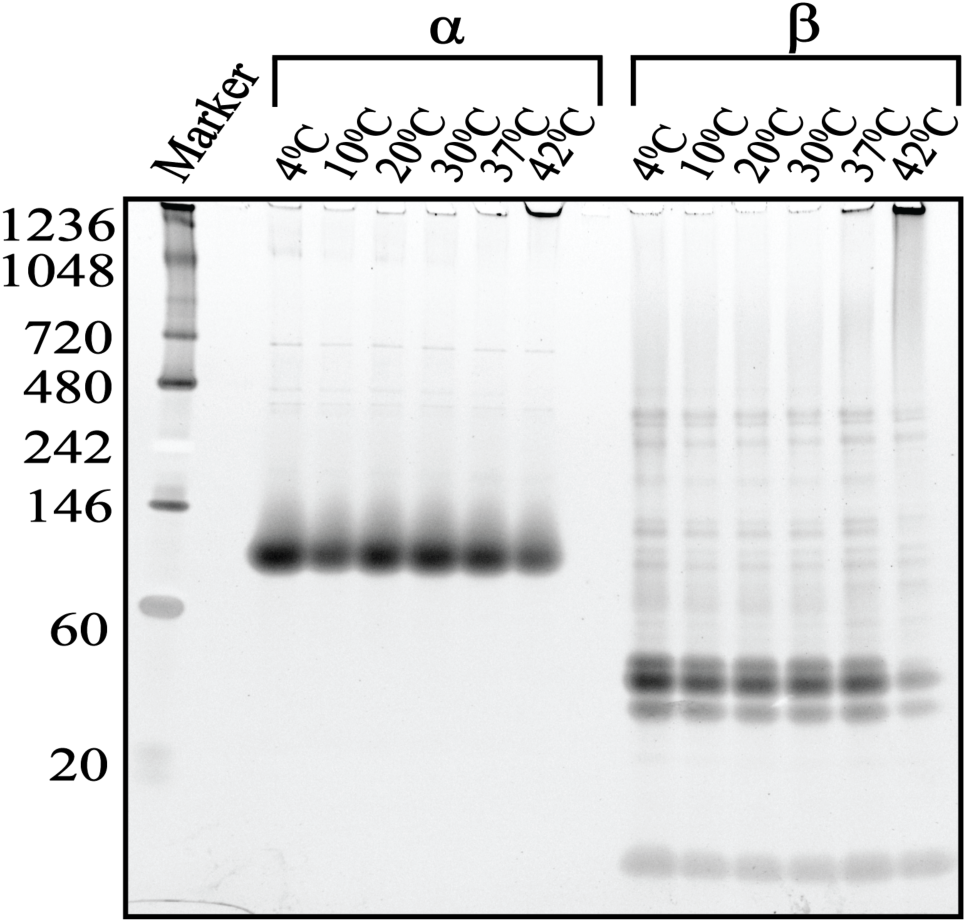
Temperature stability of the α and β subunit. 4-20% Tris Glycine native gel showing stability of the subunits over a temperature range of 4**^0^**C-42^0^C. Gel was stained with Sypro Ruby protein Stain and visualized using a Biorad ChemiDoc Imager.

### In vitro reconstitution of the *M.tb* proteasome

Having established that we have well-behaved monomeric α and β, our next goal was to study the dynamics of *M.tb* proteasome assembly. To do this we first conducted an assembly assay by mixing the α and β subunits in equimolar ratio and incubated the reaction for 24 hrs (**Fig. 4A**). All samples were loaded on a 4-20% Tris Glycine native gel, which enabled us to distinguish single subunits from the HP and full CP, as has been done previously for proteasomes from *R.e* and *Thermoplasma* (8–11). We observed a major band of 26.8 kDa for β, whereas, similar to observations made in Rhodococcus (8–11), a major band was observed around 60 kDa for alpha, consistent with these subunits remaining monomeric as we determined by DLS. Upon combining the α and β subunits, over time we observed a reduction in these individual subunits, combined with an appearance of two species of higher molecular weight. The first one to appear has an estimated MW of ∼480 kDa, consistent with the HP, consisting two stacked rings of 7α and 7β. The second species appears with a ∼2 hour delay and has an estimated MW of ∼720kDa. This is consistent with a fully assembled CP. These data are consistent with current models established in other species where two half proteasomes dimerize into a full CP.

**Fig 4:**
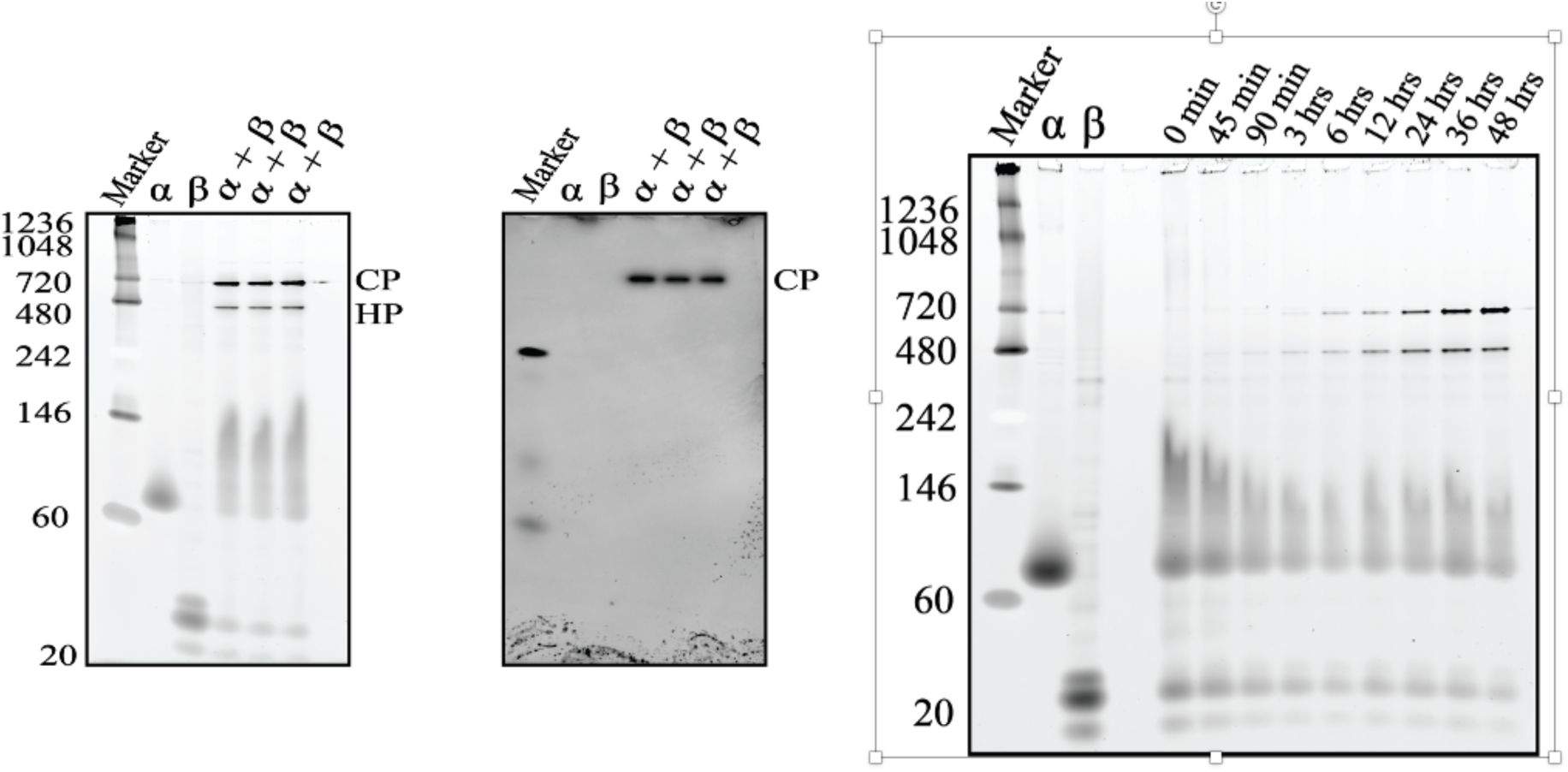
In vitro assembly of the M.tb proteasome core particle. **(A)** α and β subunits were incubated separately or together for 24 hours at 30^0^C degrees. Samples were separated on 4-20% Tris-glycine native and (A) stained with Sypro Ruby protein stain and imaged using Biorad ChemiDoc imager, or **(B)** used for In gel activity assay with 50uM Z-VLR-AMC as a substrate. Gel was imaged for activity using a 590/110 nm filter on a Biorad ChemiDoc imager. **(C)** 4-20% Tris-Glycine native gel showing the progression of assembly over the course of 2 days. Samples were collected at the time points mentioned above the gel.

To determine if the higher MW band observed in our native gels (**Fig. 4A**) was the active CP, we performed an in-gel activity assay using the fluorogenic substrare Z-VLR-AMC. The β subunit is produced as an inactive zymogen with an N-terminal pro-peptide sequence that keeps the active site sequestered. In *Rhodococcus*, the N-terminal pro-peptide is known to be autocatalytically cleaved only once two half proteasomes dimerize (8, 12). Consistent with previous findings, the beta subunit and the HP do not show proteolytic activity, but the putative CP band does (**Fig. 4B**). This confirms that the subunits self-assemble into active CPs.

### Assembly kinetics

We then performed a time-course to study the assembly of the subunits into CP over a duration of 0 mins to 2 days. The result of this time course is shown in **Fig. 4C**. We found that the individual subunits stayed monomeric over the entire time course at 4^0^C. On mixing the α and β subunits together, a smear was observed at the 0 mins point. This smear reduced over the course of the assembly experiment, suggesting that these represent smaller intermediates that assemble into the HP (**Fig. 4C**). Bands corresponding to the HP appear as early as 45mins whereas CP assembly begins only after 1.5 h of incubation and continues to increase over 48h. This indicates that both HP formation and dimerization is considerably slower as compared to *R.e,* which under the same conditions forms the HP as early as it can be measured and assembles into CP within minutes of incubation (8, 11, 12). Higher molecular weight bands were observed at the 24h, 36h and 48h time points. We do not currently know what species these bands correspond to, and leave investigation of their stoichiometry and structure to future work.

### Cross-species assembly

The genera *Rhodococcus* and *Mycobacterium* belong to the same order —Mycobacteriales, so the two species are relatively closely related, and proteasome subunits from *R.e* and *M.tb* share 65% sequence similarity. Given the evolutionary relation between the two species, we were curious to see whether the subunits have evolved to share similar interfaces and similar assembly pathways. We thus incubated subunits from *R.e* and *M.tb* were incubated together for 24h in equal concentrations and in different molar ratios to test their ability to interact with each other and spontaneously assemble into the CP. In **Fig. 5A** we see that subunits from both bacteria are compatible with each other and assemble into CPs. Note that the subunits do not have the exact same MW from the two species (especially since the *M.tb* α has an 8-residue N-terminal deletion), resulting in different apparent MW for the different compositions of the CP on the native gel. Importantly, in-gel activity assays indicate that all of the CPs that were formed are active, indicating that all of the CPs formed in these experiments were able to autocatalytically activate, regardless of composition (**Fig. 5B**).

**Fig 5:**
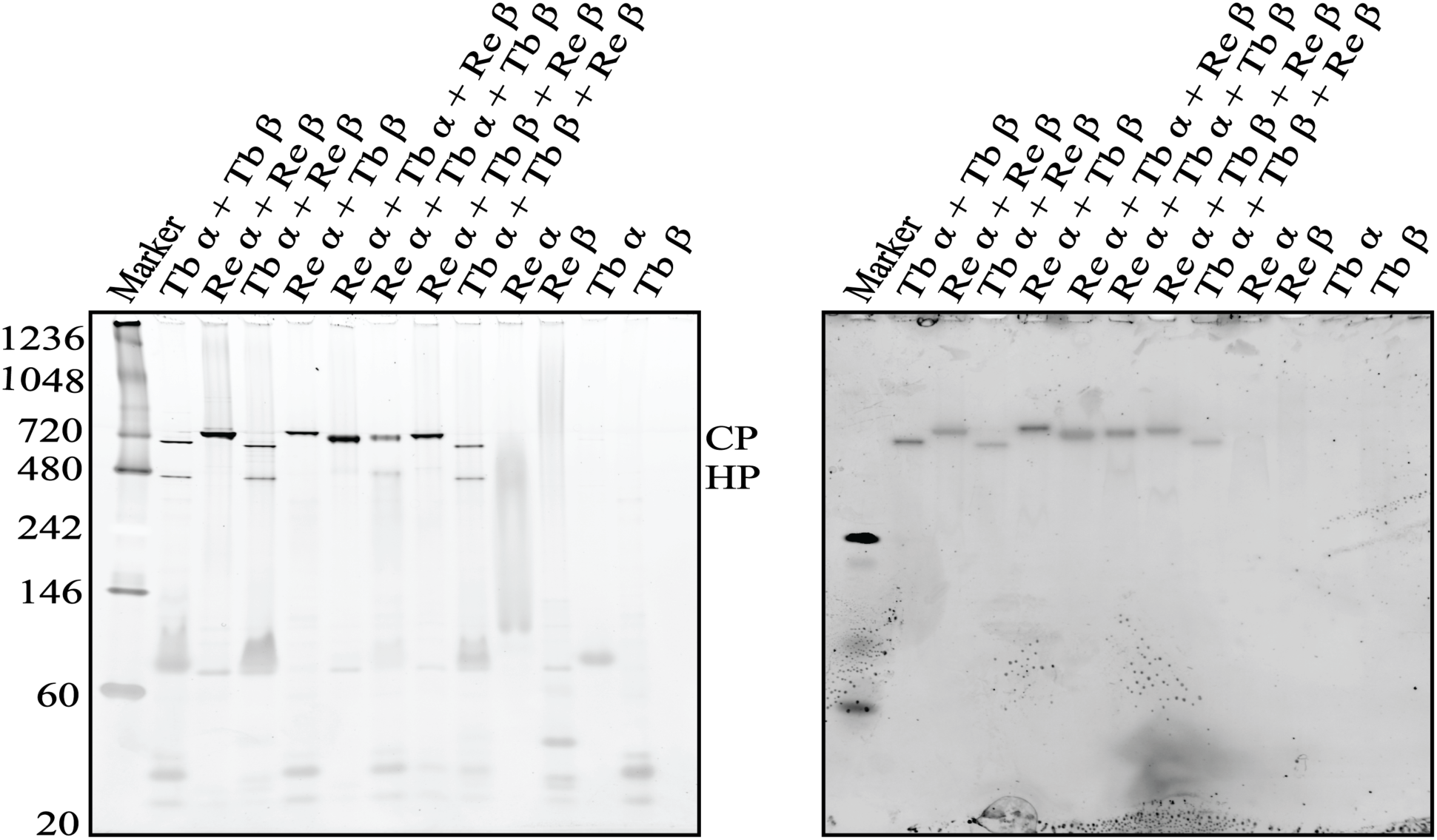
In vitro cross-assembly of R.e and M.tb proteasome subunits. **(A)** 4-20% Tris-glycine native gel showing reconstitution reactions of *R.e* and *M.tb* subunits. Ratios and composition of the subunits mentioned on top of the gel image. Gel was stained with Sypro Ruby protein stain and imaged using Biorad ChemiDoc imager. **(B)** In gel activity assay using a 4-20% Tris-glycine native gel showing reconstitution reactions of *R.e* and *M.tb* subunits. 50uM Z-VLR-AMC was used as a substrate. Flouroscence of cleaved AMC was measured using a 590/110 nm filter on a Biorad ChemiDoc imager.

Clear HP bands were observed in all cases except for when *R.e* α was incubated with *M.tb* β (**Fig. 5A**). The β subunits need to interact with each other in the HP to form the full CP, and, as described above, there is evidence that the β propeptide regulates the dimerization rate in *R.e* (*8, 11*). Thus one might expect the species of origin of the β subunit to determine properties of dimerization. However, we see that the *M.tb* α combined with different β’s all generate HPs that do not fully dimerize on the timescale of the experiment. This suggests that the α subunits play a role in regulating how quickly the HP dimerizes to form the full CP. This is surprising, and suggests some allosteric mechanism that regulates dimerization. Allostery from the α subunits to the β active sites has been observed before in yeast (14), and a similar mechanism may be operating here.

We found that the subunits did not show any species preference in reactions where α from one species was mixed with βs from both and *vice-e-versa.* For example, in the reaction containing *R.e* α + *R.e* β + *T.b* β, bands corresponding to monomeric *R.e* β or *T.b* β are not observed in excess, meaning that both the proteins are being integrated in the CP with equal probability. Taken together, these results suggest that, despite the evolutionary diversity between the two species, the subunits can assemble with each other (**Fig. 5A**), suggesting a strong conservation of both interface geometries and assembly pathways.

### Effect of subunit concentration on assembly

Multi-protein complexes are synthesized as their subunit proteins *in vivo.* The subunits have to undergo an assembly process in order to form the functional complex. Efficiency of assembly depends on both the binding affinities between the subunits and the concentration of the subunits (13). Theoretical analyses have revealed a problem in the assembly of multi-protein complexes, especially ring-like structures in particular — a phenomenon known as kinetic trapping (13). Here, subunits are kinetically stuck in stable intermediates that cannot interact with each other to form the desired tertiary structure, thereby reducing the assembly yield. We recently used a combination of computational models and experimental data to demonstrate that the proteasome CP from *R.e* exhibits extensive kinetic trapping. In particular, increasing the subunit concentration of the *R.e* CP initially increases assembly yield, but, beyond a certain critical point, assembly efficiency decreases due to kinetic trapping (unpublished work).

To determine if *M.tb* CP assembly *in vitro* is similarly sensitive to kinetic trapping, we mixed α and β in equimolar ratio at concentrations of 0.25 μM, 0.5 μM, 1μM, 2 µM, 4 µM, 8 µM and 16 µM. After 48h, two bands were observed corresponding to a full CP and a HP in all assembly reactions except for the low concentration (0.25 µM and 0.5 µM reactions, **Fig. 6A**). Next, we used these data to determine the assembly efficiency. This was calculated by measuring the CP band intensity and dividing it by individual subunit concentration (**Fig. 6B**). Assembly yield increased with increasing subunit concentrations up to around 2 μM. Further increase in subunit concentrations still yielded an overall increase in CP, however, the efficiency declined. Mathematical models described in previous studies ((13) and unpublished work) suggest that, in the low concentration (0.25µM-2µM) regime, assembly is limited by diffusion and hence increases with increase in subunit concentration. Upon further increase in concentration, these models predict that more and more subunits get kinetically trapped into stable intermediates that can cannot interact to form an HP. Dissociation of these stable intermediates occurs on very long time scales, thus depleting the monomers in the solution and reducing the assembly yield. Our results in *M.tb* are consistent with these predictions, suggesting that kinetic trapping is occurring in this case. Indeed, the assembly yield curve for *M.tb* is strikingly similar to that for *R.e* (unpublished work), again suggesting that the two CPs follow the same, evolutionarily-conserved assembly pathway. Further work, including direct observation of the kinetically trapped intermediates, will be needed to fully establish the extent and nature of kinetic trapping in these systems.

**Fig 6:**
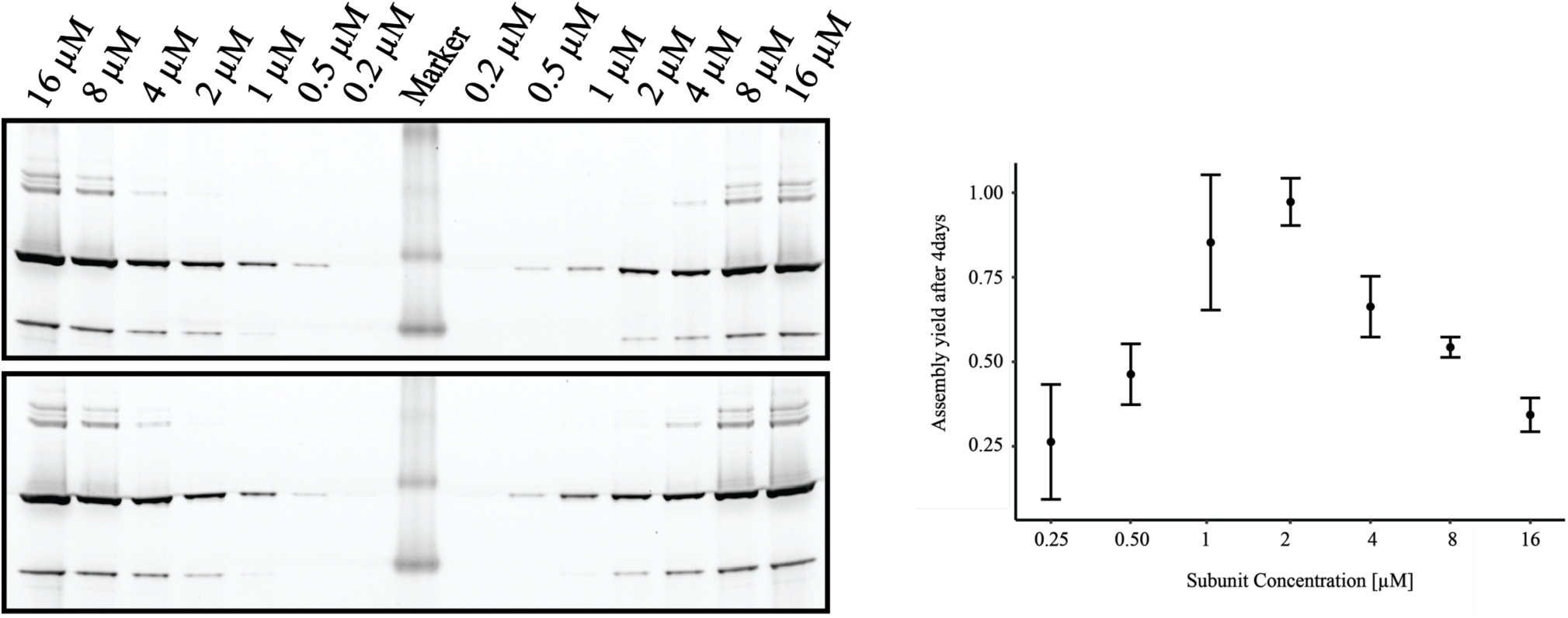
Kinetic trapping in M.tb proteasome assembly. **(A)** Representative 4-20% Tris-Glycine native gel showing CP assembly yield after 4days incubation at 30^0^C. Subunit concentrations mentioned above the gel. Gel was stained with Sypro Ruby protein satin and visualized using a Biorad ChemiDoc Imager. **(B)** quantification of gel in (A). Quantification was performed using ImageLab software. Error bars indicate standard deviation in the replicates, n=4

## Discussion

According to the WHO, drug-resistant diseases could cause 10 million deaths each year by 2050. Currently, at least 700,000 people die each year due to drug-resistant diseases, including 230,000 people who die from multi-drug resistant tuberculosis. To address this global threat of multi-drug resistance, the WHO has recommended immediate engagement from across governments, private sectors and academia. More efforts are required to develop novel antibiotics to treat infections caused by ever-evolving bacteria.

Genomic mutations confer antibiotic resistant to the bacteria but the proteasome has also been shown to play an important role in survival and pathogenicity of *M.tb*. The mammalian proteasome shares highly similar active site with that of *M.tb*, and the lack of specific enzymatic inhibitors for the proteasome suggests that targeting the proteasome active site may be difficult. In this study, we have expressed and purified the individual α and β subunits of the *M.tb* proteasome and demonstrated spontaneous *in vitro* assembly of the *M.tb* CP for the very first time. The *M.tb* subunits are stable in the temperature range of 4^0^C to 30^0^C. DLS and Native PAGE studies show that the subunits are monomeric by themselves but spontaneously assemble into active core particles on being mixed together. Assembly time course experiments indicate that the CP assembly is slow compared to *R.e*.

Interestingly, *M.tb* subunits are compatible with *R.e* subunits and on reconstitution assemble into active CPs, suggesting that proteasome subunits from the two organisms share enough structural similarity to interact with each other. The activity of the hybrid proteasomes indicates that the maturation i.e. autocatalytic cleavage of the of the N-terminal propeptide of the β subunit is not affected by the fact that these CPs are hybrids of subunits from two different species. This strongly suggests that *M.tb* and *R.e* proteasomes share a similar pathway of assembly. Furthermore, increasing the subunit concentration in *M.tb* produces a characteristic pattern of assembly efficiency almost identical to that observed in *R.e*. This is consistent with theoretical predictions of kinetic trapping ((13) and unpublished work), providing further evidence that the *M.tb* CP follows an assembly pathway shared with *R.e*, and likely conserved across all bacteria.

Establishing an *in vitro* assembly assay for the proteasome from this medically-important organism is an important step for the field because: (1) The proteasome has been established as a promising potential therapeutic target to treat M.tb infections. (2) We need drug targets to treat this disease as *M.tb* infections are harder to treat due to resistance. (3) Drugs against *M.tb* proteasomes have been difficult to develop, likely because there is substantial conservation between *M.tb* and eukaryotic proteasome active sites and targeting human proteasomes would likely cause cytotoxicity at the concentrations required to treat TB. This opens up the possibility of straightforward experimental screens for assembly inhibitors that target the *M.tb* CP, an as-of-yet completely unexplored potential approach to the treatment of TB.

In addition to providing a basis for future drug screens, the ability to perform *in vitro* assembly assays with the *M.tb* proteasome opens up other fascinating avenues of potential future research. One involves the characterization of HP assembly intermediates. In *R.e*, the HP forms so quickly that it is hard to trap intermediates for further characterization by tools like Mass Spec (15) or structural approaches like Cryo-EM. The comparatively slow assembly of the HP in *M.tb*, and the fact that we have some evidence that assembly intermediates are formed in native gels, suggests that such characterization may be possible for *M.tb*. Also, the extent of kinetic trapping in this case suggests that we may also be able to characterize kinetically trapped intermediates and compare them with those formed at lower concentration during HP assembly. Thus, in addition to establishing the conservation of assembly pathways in the CP of bacteria, our work will enable a plethora of future studies aimed at both understanding and targeting the assembly of the CP in *Mycobacterium tuberculosis*.

## Abbrevitaions

CP: Core particle
HP: Half proteasome
*R.e*: *Rhodococcus erythropolis*
*M.tb.*: *Mycobacterium tuberculosis*
Tb: tuberculosis
WHO: World Health Organization

## Notes

### Competing Interest Statement

The authors have declared no competing interest.

