## Supplementary Information for "*In vitro* reconstitution of the *M.tb* proteasome core particle reveals conserved aspects of bacterial proteasome assembly"

Amino acid sequences from PDB

**Alpha subunit (Full Length)**

MSFPYFISPEQAMRERSELARKGIARAKSVVALAYAGGVLFVAENPSRSLQKISELYDRVGFAAA

GKFNEFDNLRRGGIQFADTRGYAYDRRDVTGRQLANVYAQTLGTIFTEQAKPYEVELCVAEVAH

YGETKRPELYRITYDGSIADEPHFVVMGGTTEPIANALKESYAENASLTDALRIAVAALRAGSADT

SGGDQPTLGVASLEVAVLDANRPRRAFRRITGSALQALLVDQESPQSDGESSG

**Beta Subunit 2JAY**

MTWPLPDRLSINSLSGTPAVDLSSFTDFLRRQAPELLPASISGGAPLAGGDAQLPHGTTIVALKY

PGGVVMAGDRRSTQGNMISGRDVRKVYITDDYTATGIAGTAAVAVEFARLYAVELEHYEKLEGV

PLTFAGKINRLAIMVRGNLAAAMQGLLALPLLAGYDIHASDPQSAGRIVSFDAAGGWNIEEEGY

QAVGSGSLFAKSSMKKLYSQVTDGDSGLRVAVEALYDAADDDSATGGPDLVRGIFPTAVIIDADG

AVDVPESRIAELARAIIESRSGADTFGSDGGEK

**<u>Gene Sequences</u>**

**PrcA gene**

NNNNNNNNNNNNNCNNNNNNGTGCCGGCTCCGGAGAGCTCTTTAATTAAGCGGCCGCCCT

GCAGGACTCGAGTTCTAGAAATAATTTTGTTTAACTTTAAGAAGGAGATATACATATGAAATCTT

CTCACCATCACCATCACCATGAAAACCTGTACTTCCAATCCAATGCAATGGAGCAGGCGATG

CGTGAGCGTAGCGAACTGGCGCGTAAGGGTATCGCGCGTGCGAAAAGCGTGGTTGCGCTG

GCGTACGCGGGTGGCGTGCTGTTCGTTGCGGAGAACCCGAGCCGTAGCCTGCAGAAGATCA

GCGAACTGTATGACCGTGTGGGTTTCGCGGCGGCGGGCAAATTCAACGAATTTGACAACCTG

CGTCGTGGTGGCATTCAATTTGCGGATACCCGTGGTTACGCGTATGACCGTCGTGATGTGAC

CGGCCGTCAGCTGGCGAACGTTTACGCGCAAACCCTGGGCACCATTTTCACCGAGCAAGCG

AAGCCGTACGAGGTTGAACTGTGCGTGGCGGAAGTTGCGCACTATGGCGAGACCAAACGTC

CGGAACTGTACCGTATCACCTATGACGGCAGCATTGCGGATGAGCCGCACTTTGTGGTTATG

GGTGGCACCACCGAACCGATCGCGAACGCGCTGAAGGAGAGCTATGCGGAAAACGCGAGC

CTGACCGATGCGCTGCGTATTGCGGTGGCGGCGCTGCGTGCGGGTAGCGCGGACACCAGC

GGTGGCGATCAGCCGACCCTGGGCGTTGCGAGCCTGGAAGTGGCGGTTCTGGATGCGAAC

CGTCCGCGTCGTGCGTTTCGTCGTATTACCGGTAGCGCGCTGCAGGCGCTGCTGGTTGACC

AGGAGAGCCCGCAAAGCGATGGTGAAAGCAGCGGCTAATAATAACATTGGAAGTGGATAACG

GATCCGCGATCGCGGCGCGCCACCTGGTGGCCGGCCGGTACCACGCGTGCGCGCTGATCC

GGCTGCTAACAAAGCCCGAAGGAAGCTGAGTTGGCTGCTGCCACCGCTGAGCATAACTAGC

ATANCCCTTGGGGCNTCTAACGGGTCTGAGGGTTTTTGCTGAANGNAGGAACTANNTCCGGA

NNTCNNNNGGANGGNGNNGNCNCCATGATCGCGTANTCNNTNGTGNNNCAGTANCNANCN

NNNNGNNNGNNNNNNNNAGCGNNGNNNNNNNNNCNANNANGGGTGNCATNNAANTGNNNNN

ANNCNTNTNNNN NNNNN

<u>Amino Acid Sequence translated from above code</u>

MKSSHHHHHHENLYFQSNAMEQAMRERSELARKGIARAKSVVALAYAGGVLFVAENPSRSLQKI

SELYDRVGFAAAGKFNEFDNLRRGGIQFADTRGYAYDRRDVTGRQLANVYAQTLGTIFTEQAKPY

EVELCVAEVAHYGETKRPELYRITYDGSIADEPHFVVMGGTTEPIANALKESYAENASLTDALRIAVA

AL RAGSADTSGGDQPTLGVASLEVAVLDANRPRRAFRRITGSALQALLVDQESPQSDGESSG—H

NOTE - First 8 amino acids were deleted from the sequence to make it an open gate variant

**Prc B gene**

NNNNNNNNNTTNNNNCTAGTGCCGGCTCCGGAGAGCTCTTTAATTAAGCGGCCGCCCTGCA

GGACTCGAGTTCTAGAAATAATTTTGTTTAACTTTAAGAAGGAGATATAGATCATGACCTGGCC

GCTGCCGGATCGTCTGAGCATTAACAGCCTGAGCGGTACCCCGGCGGTTGATCTGAGCAGC

TTCACCGACTTTCTGCGTCGTCAGGCGCCGGAACTGCTGCCGGCGAGCATTAGCGGTGGCG

CGCCGCTGGCGGGTGGCGACGCGCAACTGCCGCACGGTACCACCATCGTGGCGCTGAAGT

ATCCGGGTGGCGTGGTTATGGCGGGCGATCGTCGTAGCACCCAGGGTAACATGATCAGCGG

CCGTGACGTGCGTAAAGTTTACATCACCGACGATTATACCGCGACCGGTATTGCGGGTACCG

CGGCGGTGGCGGTTGAGTTCGCGCGTCTGTACGCGGTGGAACTGGAGCACTATGAAAAGCT

GGAGGGTGTTCCGCTGACCTTTGCGGGCAAAATCAACCGTCTGGCGATTATGGTTCGTGGTA

ACCTGGCGGCGGCGATGCAAGGTCTGCTGGCGCTGCCGCTGCTGGCGGGTTACGATATTCA

TGCGAGCGACCCGCAAAGCGCGGGTCGTATTGTGAGCTTCGATGCGGCGGGTGGCTGGAA

CATCGAGGAAGAGGGTTACCAGGCGGTTGGTAGCGGCAGCCTGTTTGCGAAGAGCAGCATG

AAGAAACTGTATAGCCAAGTGACCGATGGTGACAGCGGTCTGCGTGTGGCGGTTGAAGCGCT

GTATGATGGGCGGATGATGATAGCGCGACCGGTGGCCCGGACCTGGTGCGTGGTATTTTCC

CGACCGCGGTTATCATTGATGCGGATGGTGCGGTGGATGTTCCGGAAAGCCGTATCGCGGA

GCTGGCGCGTGCGATCATTGAAAGCCGTAGCGGTGCGGATACCTTTGGCAGCGANGNGGC

GAGAAACACCACCNCCACCACCACTNATAATAAGATCCCNACTCANTAAGGATCCGCGATCG

CGGNGCGCCANCTGGNNGNNGGNNGTACCANNCGTGCNCGCTGATCCGGCTGCTAACAAA

NNCCCGAAAGGAAGCTGANTTGGCTGCTGCCNNCGCTGNNNATNACTANCNNNNCCCNTGG

GGNNNNNAAAC GGTNNTGAGGNNNTTTTNNNNNNNAAGGNNNNGNNN

Amino Acid Sequence translated from above code

MTWPLPDRLSINSLSGTPAVDLSSFTDFLRRQAPELLPASISGGAPLAGGDAQLPHGTTIVALKYP

GGVVMAGDRRSTQGNMISGRDVRKVYITDDYTATGIAGTAAVAVEFARLYAVELEHYEKLEGVPLT

FAGKINRLAIMVRGNLAAAMQGLLALPLLAGYDIHASDPQSAGRIVSFDAAGGWNIEEEGYQAVGS

GSLFAKSSMKKLYSQVTDGDSGLRVAVEALYDAADDDSATGGPDLVRGIFPTAVIIDADGAVDVPE

SRIAELARAIIESRSGADTFGSXXARNTTXTTTXNKIXTX

***Sup. Table 1:Plasmids, primers and dNTPs used in cloning the M.tb PrcA and Prc B genes***

| **Gene** | **Vector** | **Forward primer 5'-3'** | **Reverse primer 5'-3'** | **dNTP for Insert** | **dNTP for Vector** |
|---|---|---|---|---|---|
| Prc A | 2B-T | TACTTCCAATCCAATGCAATGGAGCAAGGCGATGCG | TTATCCACTTCCAATGTTATTATTAGCCGCTGCTTTCACC | G | C |
| Prc B | 2A-T | TTTAAGAAGGAGATATAGATCATGCATCATCATCATCACCA | TTATGGAGTTGGGATCTTATTATTAGTGGTGGTGGTGGT | C | G |

### *Expression and purification of the Rhodococcus erythrpolis α1 subunit*

The α1 gene of *R. erythropolis* was subcloned from a pT7 plasmid (a generous gift from Wolfgang Baumeister) in a pTBSG plasmid background with a N terminal 6xHis-tag followed by a tobacco etch virus(TEV) protease cleavage site by Philip Gao and Anne Cooper at the Protein Production core lab, University of Kansas. Plasmids were freshly transformed in *E. coli* BL21 (DE3)-pLyse S(NEB) cells and were grown in 4 Liters Luria Barteni (LB) media containing 100 μg/ml Carbenicillin and 30 μg/ml

Chloramohenicol. Bacteria were grown in a shaking incubator at 200rpm and 37°C, induced with 1 mM isopropyl-β-D-thiogalactopyranoside (IPTG) at OD600 ~ 0.6 and cells were allowed to grow overnight at 15°C. Cells were harvested by centrifugation at 4000rpm for 12 mins using a (JS-4.750 rotor in a Avanti-J15R centrifuge), resuspended in lysis buffer (50mM Tris, 100mM NaCl, 10mM MgCl2, 1mMATP, 5mM BME, 30% glycerol pH7.05 at RT), and sonicated at 45% amplitude with 2s on - 10s off cycles for 4 mins a using a Fischer Scientific Sonic dismembrator model 500. Cellular debris were removed by centrifugation at 10,000rpm for 30 min in a JA-10.100 rotor in a Avanti-J15R centrifuge. Clarified lysate was passed through a 10 ml $Ni^{2+}$-Sepharose 6 Fast flow resin (GE Healthcare Catalogue number 17-5318-02), washed with 50 ml of binding buffer (50mM Tris, 100mM NaCl, 10mM MgCl2, 1mMATP, 5mM BME, 30% glycerol pH7.05 at RT) followed by a second wash with 50ml wash buffer (50mM Tris, 100mM NaCl, 10mM MgCl2, 1mM ATP, 5mM BME, 30% glycerol, 10mM Imidazole pH7.05 at RT) and eluted with a total of 50 ml elution buffer (50mM Tris, 100mM NaCl, 10mM MgCl2, 1mMATP, 5mMBME, 30% glycerol, 1M Imidazole pH7.05 at room temperature). All fractions were analyszed using SDS-PAGE. Fractions containing purified protein were incubated with 1:20 v/v of 62 μM TEV protease(1ml TEV/20mL protein) and dialyzed overnight in binding buffer. Dialyzed proteins were subjected to the exact same $Ni^{2+}$-affinity chromatography to separate the protein from cleaved 6xHis-tags. Purified protein was then mixed with 20% v/v glycerol, aliquoted in 1ml vials and stored in -80°C for later use.

#### *Expression and purification of the Rhodococcus erythropolis β1 subunit*

Initial attempts of expressing and purifying the β subunit from pT-7 and pTBSG plasmids (generous gifts from Wolfgang Baumeister and Philip Gao respectively) failed to produce pure protein. Hence, DNA corresponding to the β1 gene of *R. erythropolis* optimized for expression in *E.coli* was purchased from Genscript. The optimized gene was PCR amplified and subcloned into the NdeI/XhoI sites of pET-22b (a generous gift of Roberto DeGuzman, University of Kansas), which introduced a C-terminal His6-tag for protein purification to generate the pAKRE-β plasmid. Plasmid was freshly transformed in *E. coli* BL21

(DE3) and cells were grown in 4L LB media containing 100 μg/ml carbenicillin. Bacteria were grown in a shaking incubator at 200rpm and 37°C, induced with 1mM isopropyl-β-D-thiogalactopyranoside (IPTG) at OD600 ~ 0.6 and cell growth was continued for 4hours at 30°C. Cells were harvested by centrifugation at 4000rpm for 12 mins using a JS-4.750 rotor in a Avanti-J15R centrifuge and cell pellets were resuspended in 50 ml binding buffer (25mM Na2PO4, 300mM NaCl, 5mM Imidazole pH7.4). Cells were lysed by sonicating at 45% amplitude with 2s on-10s off cycles for 4 mins. Cellular debris were removed by centrifugation at 10,000rpm for 30 min in a JA-10.100 rotor in a Avanti-J15R centrifuge. Following centrifugation, the protein was purified using a GE AKTA pure FPLC system. The cleared lysate was loaded on a Nickel affinity column (GE His-trap 5ml column), the column was then washed with 10 column volumes (CV) of binding buffer (25mM Na2PO4, 300mM NaCl, 5mM Imidazole pH7.4). The bound protein was then eluted with a linear gradient of the elution buffer (25mM Na2PO4, 300mM NaCl, 500mM Imidazole pH7.4) over 20CV. Peak fractions were analyzed using SDS-PAGE. Fractions containing the protein were dialyzed in anion exchange binding buffer (20mM Tris-HCL, 20mM NaCl, pH 8.0 at $4^0$C) overnight at $4^0$C. On the second day, dialyzed protein was cleaned up using a GE 5ml Hi-trap anion exchange column. Unbound protein and contaminants were washed with 10 CV of binding buffer (20mM Tris-HCL, 20mM NaCl, pH 8.0 at $4^0$C). Bound protein was eluted using a linear gradient of the elution buffer (20mM Tris-HCL, 1M NaCl, pH 8.0 at $4^0$C) over 20CV. Peak fractions were analyzed using SDS-PAGE. Fractions containing the protein were pooled together. 20% V/V glycerol was added to the pooled fractions and aliquotes of 300μL were frozen in liquid nitrogen and stored at $-80^0$C for later use

Raw data for **Fig.7B**

| Concentration | CP I 1 | AY1 | CP I 2 | AY2 | CP I 3 | AY3 | CP I 4 | AY4 | Assembly Yield | STD | Normalized AY1 | Normalized AY2 | Normalized AY3 | Normalized AY4 | Norm. Av. AY | Std. of.norm.AY |
|---|---|---|---|---|---|---|---|---|---|---|---|---|---|---|---|---|
| 0.25 | 4.0E+04 | 1.6E+05 | 5.6E+04 | 2.2E+05 | 1.7E+05 | 6.6E+05 | 5.0E+04 | 2.0E+05 | 3.1E+05 | 2.4E+05 | 0.15 | 0.17 | 0.50 | 0.18 | 0.25 | 0.17 |
| 0.5 | 2.3E+05 | 4.5E+05 | 3.5E+05 | 6.9E+05 | 3.2E+05 | 6.4E+05 | 1.9E+05 | 3.8E+05 | 5.4E+05 | 1.5E+05 | 0.44 | 0.54 | 0.48 | 0.33 | 0.45 | 0.09 |
| 1 | 5.7E+05 | 5.7E+05 | 1.2E+06 | 1.2E+06 | 1.3E+06 | 1.3E+06 | 9.8E+05 | 9.8E+05 | 1.0E+06 | 3.4E+05 | 0.55 | 0.96 | 1.00 | 0.86 | 0.84 | 0.20 |
| 2 | 2.1E+06 | 1.0E+06 | 2.6E+06 | 1.3E+06 | 2.3E+06 | 1.1E+06 | 2.3E+06 | 1.1E+06 | 1.1E+06 | 1.0E+05 | 1.00 | 1.00 | 0.85 | 1.00 | 0.96 | 0.07 |
| 4 | 2.4E+06 | 5.9E+05 | 3.5E+06 | 8.8E+05 | 3.1E+06 | 7.7E+05 | 3.5E+06 | 8.7E+05 | 7.8E+05 | 1.3E+05 | 0.57 | 0.68 | 0.58 | 0.77 | 0.65 | 0.09 |
| 8 | 4.4E+06 | 5.5E+05 | 5.3E+06 | 6.6E+05 | 5.2E+06 | 6.5E+05 | 5.1E+06 | 6.4E+05 | 6.2E+05 | 5.1E+04 | 0.53 | 0.51 | 0.49 | 0.56 | 0.53 | 0.03 |
| 16 | 4.7E+06 | 2.9E+05 | 6.6E+06 | 4.1E+05 | 6.4E+06 | 4.0E+05 | 7.2E+06 | 4.5E+05 | 3.9E+05 | 6.8E+04 | 0.29 | 0.32 | 0.30 | 0.40 | 0.33 | 0.05 |
